# A reference database of reptile images

**DOI:** 10.1101/2024.03.08.584020

**Authors:** Peter Uetz, Maya Patel, Zainab Gbadamosi, Jeremy Nguyen, Stacey Shoope

## Abstract

While there are millions of reptile images available online, they are not well organized and not easily findable, accessible, interoperable, or reproducible (FAIR). More importantly, they are not standardized and thus hardly comparable. Here we present a reference database of more than 14,000 standardized images of 1,045 reptile species (969 lizard and 76 snake species), that are based on preserved specimens in 20 different collections, including 533 type species of genera and type specimens of 72 species. All images were taken with standardized views, including dorsal and ventral body shots as well as dorsal, ventral, and lateral views of the heads and other body parts. Although only 11 out of the 20 collections are cross-referenced in Vertnet, some others are indexed in GBIF, and this fraction will certainly grow in the near future. The utility of this and similar image collections will further grow with additional material and further cross-referencing, e.g., to DNA sequence databases or citizen science projects. The images are searchable and freely available on Morphobank.org (Project 5121) and on Figshare.com.

## Introduction

Despite the critical importance of images in biomedicine, there are relatively few image databases that provide structured, annotated, and standardized collections of images using FAIR guidelines, i.e., they are findable, accessible, interoperable, and reusable. For images related to biodiversity and taxonomy, two such databases are Morphobank (O’Leary & Kaufman 2011) and Morphosource (www.morphosource.org). These repositories focus on collecting images and traits in a phylogenetic context (Morphobank) and on 3D images such as CT scans (Morphosource), respectively.

There is no shortage of animal photos on the web and several attempts have been made to survey them in a more systematic way, including reptiles (Durso et al. 2021; Marshall et al. 2020). Some databases such as iNaturalist have huge image collections counting in the millions but their main purpose is to document observations in nature. Although iNaturalist is not a taxonomic project, it has recently started to add images of museum specimens. Many taxon-specific databases, such as the Reptile Database, focus on taxonomy and but not specifically on image collection. We believe that taxonomists need specialized databases that provide standardized images, e.g., in order to permit direct comparison of specimens in high resolution so that diagnostic features can be easily recognized. While many natural history collections offer images on their websites, this is only true for a subset of collections or a subset of their specimens. Importantly, the distributed nature of having images spread over numerous websites makes those images hard to find and even harder to use.

The purpose of this project is to provide a starting point for a systematic collection of reptile images, taken from preserved specimens that can be revisited in those collections, with the pertinent meta-data such as localities and other information. We started by taking pictures of more than 1,000 reptile species in 20 collections around the world, resulting in more than 14,000 images that have been tagged with meta-data, so that they can be sorted by specific features such as particular body parts and cross-referenced with other databases such as VertNet or the Reptile Database. We are making this collection accessible through Morphobank, where each image can be traced back to a specific specimen based on its collection and catalogue number.

As an example and case study, we present photos of 8 species of the subfamily Lacertinae (currently identical with the tribus Lacertini), an Old World group containing 19 genera and 139 species of lizards to show the utility of such image collections for comparative studies (Fig. 1).

**Fig. 1.**
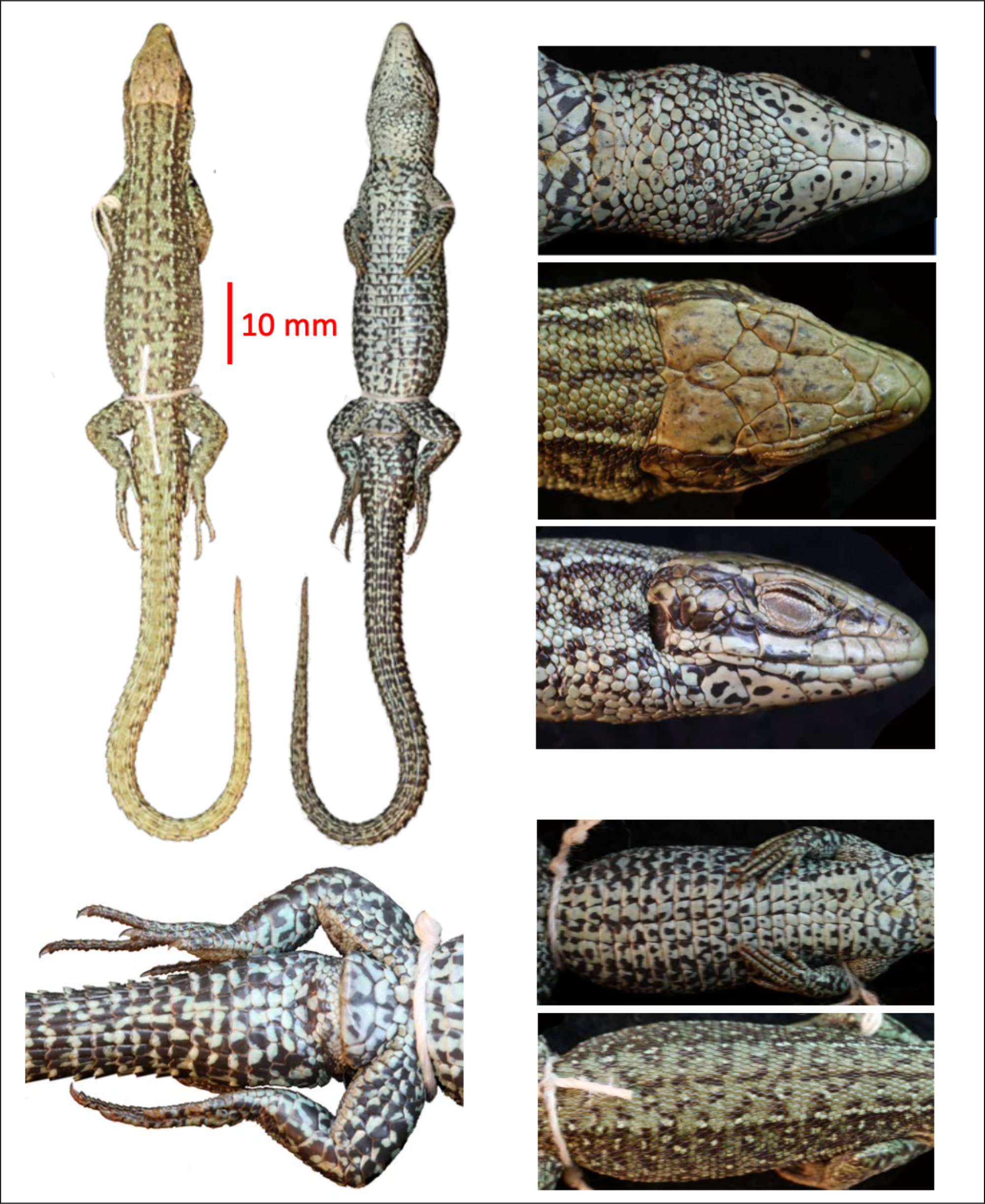
Sample images, showing different views of one specimen, ZSM 2003/2006, representing *Zootoca vivipara* Lichtenstein 1823. Almost all specimens in the image collection are shown as the whole animal with dorsal and ventral views and with dorsal, lateral, and ventral head shots. In addition, the trunk is usually shown from dorsal and ventral views, and many specimens show additional features such as the anal area. All specimens include rulers for size measurements.

## Methods and Materials

A total of 62.6 GB of 15,675 photographs encompassing 1,045 reptile species were taken across 20 collections around the world (**Table 1**). 14,239 of these photos show specimens, the remaining 1436 show labels which may or may not contain additional information such as localities. The photos depict 1,342 unique specimens, with the explicit intention to take pictures from the same perspective, namely the dorsal and ventral side of the whole body, plus close-ups of the head (dorsal, lateral, and ventral sides), as well as details of the trunk (lateral view) and the cloacal region. In some cases, close-ups of other body parts were taken too (such as the toe pads in geckos), depending on preservation state and other factors.

**Table 1.** Collections used for imaging. For specimen and species details see Table 2.

| Collection | Full Name | Location |
| --- | --- | --- |
| AMS | Australian Museum | Sydney |
| CMNAR | Canadian Museum of Nature (herps) | Ottawa |
| MHNG | Natural History Museum of Geneva | Geneva |
| MNHN | Museum National d'Histoire Naturelle | Paris |
| NCSM | North Carolina Museum of Natural Sciences | Durham |
| NMBA | Naturhistorisches Museum Basel | Basel |
| NMV | Museum Victoria | Melbourne |
| NMW | Naturhistorisches Museum Wien | Vienna |
| QM | Queensland Museum | Brisbane |
| RMCA | Royal Museum of Central Africa | Brussels |
| SAMA | South Australian Museum | Adelaide |
| SMF | Senckenberg Naturmuseum Frankfurt | Frankfurt |
| SMNS | State Museum of Natural History Stuttgart | Stuttgart |
| TAU | Tel Aviv University | Tel Aviv |
| UF | University of Florida | Gainesville |
| WAM | Western Australian Museum | Perth |
| ZFMK | Zoologisches Forschungsmuseum Alexander Koenig | Bonn |
| ZMB | Museum für Naturkunde Berlin | Berlin |
| ZRC | Zoological Reference Collection | Singapore |
| ZSM | Zoologische Staatssammlung München | Munich |

**Table 2.** Breakdown of images by collection. Zeros indicate that none of our species is recorded from a given collection in Vertnet. N/A indicates that the collection is not represented in VertNet.

| Collection | Specimens | Species | Species in Vertnet | Species with localities |
| --- | --- | --- | --- | --- |
| AMS | 72 | 59 | 48 | 36 |
| CMNAR | 35 | 33 | 33 | 5 |
| MHNG | 24 | 19 | ? | 0 |
| MNHN | 78 | 73 | 24 | 0 |
| NCSM | 37 | 35 | 16 | 12 |
| NMBA | 85 | 65 | N/A | N/A |
| NMV | 39 | 35 | N/A | N/A |
| NMW | 99 | 49 | N/A | N/A |
| QM | 109 | 65 | 50 | 35 |
| RMCA | 63 | 54 | ? | 0 |
| SAMA | 29 | 26 | 19 | 3 |
| SMF | 64 | 63 | N/A | N/A |
| SMNS | 34 | 29 | 13 | 3 |
| TAU | 17 | 13 | N/A | N/A |
| UF | 113 | 98 | ? | 0 |
| WAM | 73 | 48 | 48 | 34 |
| ZFMK | 190 | 131 | N/A | N/A |
| ZMB | 97 | 88 | N/A | N/A |
| ZRC | 35 | 33 | N/A | N/A |
| ZSM | 51 | 49 | N/A | N/A |

Since many photos were similar and created some redundancy, we removed some of this redundancy using Duplicate Photo Cleaner version 7.5.0.12 (https://www.easyduplicatefinder.com/). This removed photos that exhibited at least an 85% similarity threshold to one or more other photos.

Metadata was added to the photos using a custom-made tool built in Claris FileMaker Pro (https://www.claris.com/filemaker/pro/). The view and visible body parts were marked for each image. Keywords for view included *dorsal, ventral, left lateral, right lateral*, and *lateral*, while keywords for body parts consisted of the *whole body, head, trunk, forelimb, hindlimb, tail, hand, foot*, and *cloaca*. After all the tagging was completed, the metadata was exported from FileMaker in the form of Excel sheets showing file names, body parts, and views for each image file.

The full-sized images and the metadata Excel sheets were available in Morphobank (https://morphobank.org) under project number 5121 and in Figshare (https://figshare.com).

## Results

We have taken more than 14,239 standardized photos representing 1,045 species of reptiles and made them available in Morphobank. These photos can serve as a reference dataset for comparative morphology, species identification, or other purposes. While only 81 of the 1,342 specimens are primary types, images of other type specimen can be added as they become available.

Taxonomically, our initial goal was to obtain photos of all lizard genera, that is, at least one species of each genus (ideally the type species). While we have not reached that goal yet, our collection has photos of 490 lizard genera (out of 603), that is, 81% of all lizard genera. However, representing all genera is a moving target: since the turn of the century (2000), 122 new lizard genera have been described (Uetz et al. 2024), including 16 genera in the years 2021 and 2022, not counting those from taxonomic vandalism (Wüster et al. 2021). Most of these were not covered by this project, given that our current set of photos were mostly taken from 2019 to 2020 (before Covid made museum visits rather difficult).

Overall, we collected photos of 53 families, 29 of which having all genera covered (Table 3). While most of these families are rather small, two of them have 10 or more genera, namely Anguidae and Phyllodactylidae. Of 41 families or nearly half of all families, we have captured 80% or more of all genera.

**Table 3.**
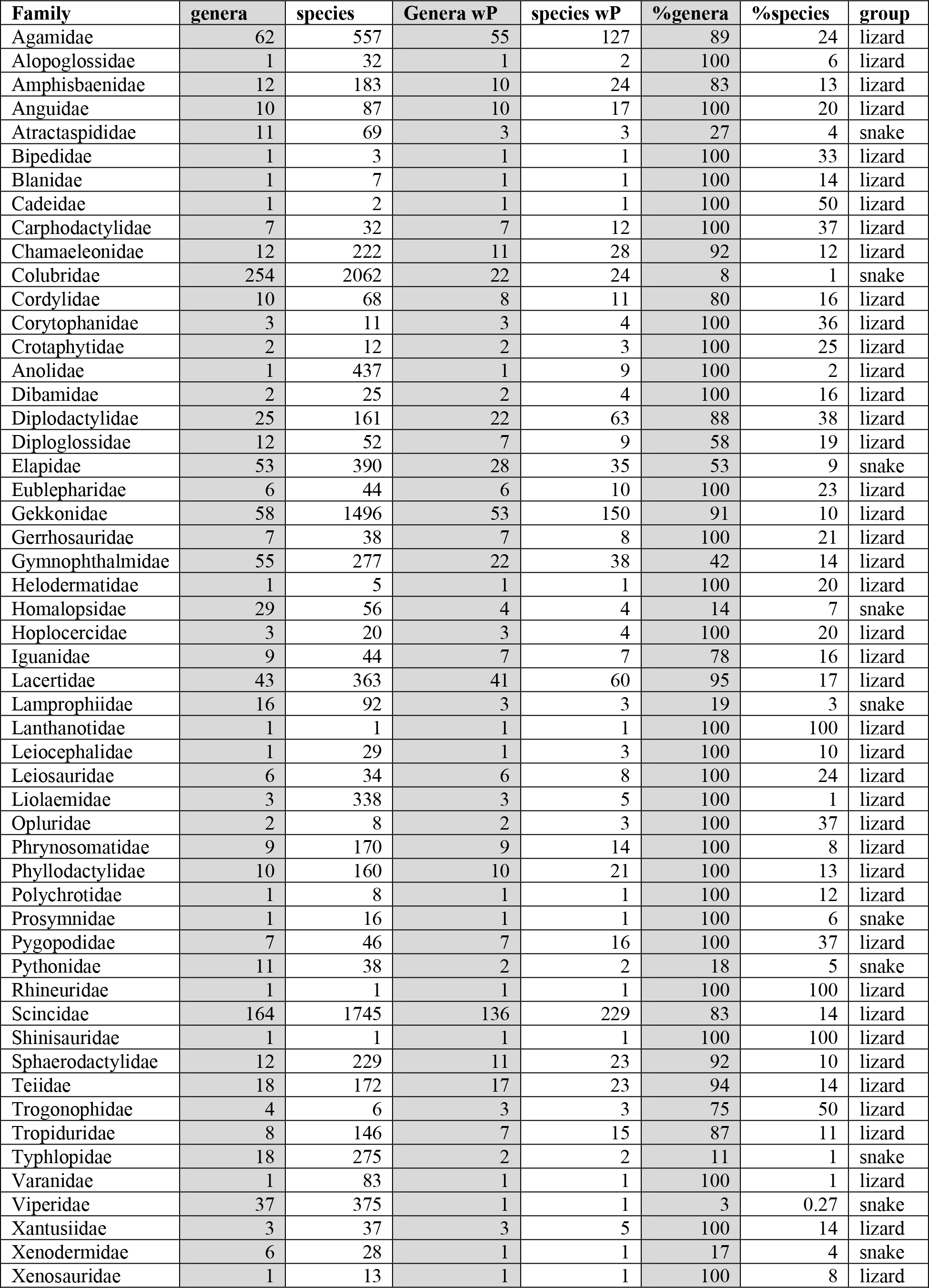
(next page). **Number of genera and species in each family with images in our collections**. wP = with photos. Numbers based on Reptile Database as of January 2024. Percentages were rounded to nearest integer.

| **Family** | **genera** | **species** | **Genera wP** | **species wP** | **%genera** | **%species** | **group** |
| --- | --- | --- | --- | --- | --- | --- | --- |
| Agamidae | 62 | 557 | 55 | 127 | 89 | 24 | lizard |
| Alopoglossidae | 1 | 32 | 1 | 2 | 100 | 6 | lizard |
| Amphisbaenidae | 12 | 183 | 10 | 24 | 83 | 13 | lizard |
| Anguidae | 10 | 87 | 10 | 17 | 100 | 20 | lizard |
| Atractaspididae | 11 | 69 | 3 | 3 | 27 | 4 | snake |
| Bipedidae | 1 | 3 | 1 | 1 | 100 | 33 | lizard |
| Blanidae | 1 | 7 | 1 | 1 | 100 | 14 | lizard |
| Cadeidae | 1 | 2 | 1 | 1 | 100 | 50 | lizard |
| Carphodactylidae | 7 | 32 | 7 | 12 | 100 | 37 | lizard |
| Chamaeleonidae | 12 | 222 | 11 | 28 | 92 | 12 | lizard |
| Colubridae | 254 | 2062 | 22 | 24 | 8 | 1 | snake |
| Cordylidae | 10 | 68 | 8 | 11 | 80 | 16 | lizard |
| Corytophanidae | 3 | 11 | 3 | 4 | 100 | 36 | lizard |
| Crotaphytidae | 2 | 12 | 2 | 3 | 100 | 25 | lizard |
| Anolidae | 1 | 437 | 1 | 9 | 100 | 2 | lizard |
| Dibamidae | 2 | 25 | 2 | 4 | 100 | 16 | lizard |
| Diplodactylidae | 25 | 161 | 22 | 63 | 88 | 38 | lizard |
| Diploglossidae | 12 | 52 | 7 | 9 | 58 | 19 | lizard |
| Elapidae | 53 | 390 | 28 | 35 | 53 | 9 | snake |
| Eublepharidae | 6 | 44 | 6 | 10 | 100 | 23 | lizard |
| Gekkonidae | 58 | 1496 | 53 | 150 | 91 | 10 | lizard |
| Gerrhosauridae | 7 | 38 | 7 | 8 | 100 | 21 | lizard |
| Gymnophthalmidae | 55 | 277 | 22 | 38 | 42 | 14 | lizard |
| Helodermatidae | 1 | 5 | 1 | 1 | 100 | 20 | lizard |
| Homalopsidae | 29 | 56 | 4 | 4 | 14 | 7 | snake |
| Hoplocercidae | 3 | 20 | 3 | 4 | 100 | 20 | lizard |
| Iguanidae | 9 | 44 | 7 | 7 | 78 | 16 | lizard |
| Lacertidae | 43 | 363 | 41 | 60 | 95 | 17 | lizard |
| Lamprophiidae | 16 | 92 | 3 | 3 | 19 | 3 | snake |
| Lanthanotidae | 1 | 1 | 1 | 1 | 100 | 100 | lizard |
| Leiocephalidae | 1 | 29 | 1 | 3 | 100 | 10 | lizard |
| Leiosauridae | 6 | 34 | 6 | 8 | 100 | 24 | lizard |
| Liolaemidae | 3 | 338 | 3 | 5 | 100 | 1 | lizard |
| Opluridae | 2 | 8 | 2 | 3 | 100 | 37 | lizard |
| Phrynosomatidae | 9 | 170 | 9 | 14 | 100 | 8 | lizard |
| Phyllodactylidae | 10 | 160 | 10 | 21 | 100 | 13 | lizard |
| Polychrotidae | 1 | 8 | 1 | 1 | 100 | 12 | lizard |
| Prosymnidae | 1 | 16 | 1 | 1 | 100 | 6 | snake |
| Pygopodidae | 7 | 46 | 7 | 16 | 100 | 37 | lizard |
| Pythonidae | 11 | 38 | 2 | 2 | 18 | 5 | snake |
| Rhineuridae | 1 | 1 | 1 | 1 | 100 | 100 | lizard |
| Scincidae | 164 | 1745 | 136 | 229 | 83 | 14 | lizard |
| Shinisauridae | 1 | 1 | 1 | 1 | 100 | 100 | lizard |
| Sphaerodactylidae | 12 | 229 | 11 | 23 | 92 | 10 | lizard |
| Teiidae | 18 | 172 | 17 | 23 | 94 | 14 | lizard |
| Trogonophidae | 4 | 6 | 3 | 3 | 75 | 50 | lizard |
| Tropiduridae | 8 | 146 | 7 | 15 | 87 | 11 | lizard |
| Typhlopidae | 18 | 275 | 2 | 2 | 11 | 1 | snake |
| Varanidae | 1 | 83 | 1 | 1 | 100 | 1 | lizard |
| Viperidae | 37 | 375 | 1 | 1 | 3 | 0.27 | snake |
| Xantusiidae | 3 | 37 | 3 | 5 | 100 | 14 | lizard |
| Xenodermidae | 6 | 28 | 1 | 1 | 17 | 4 | snake |
| Xenosauridae | 1 | 13 | 1 | 1 | 100 | 8 | lizard |

Given that our focus was on lizard genera, we attempted to obtain photos of the type species of each genus. This was achieved for 530 genera, including 478 lizard genera and for 52 snake genera (**Supplementary Table S1**).

In order to make the images more accessible, we removed about 20% of images as duplicates using a 85% similarity threshold (see methods). The remaining 14,239 image files were renamed and marked with metadata, so that they can be searched for features such as “head” or “ventral” sides. The tagged photos can be searched on or downloaded from the Morphobank and Figshare web sites (see Materials).

## Discussion

This project aims to fill the need for a reference database of specimen photos that are highly standardized in terms of what and how they show it. The image collection presented here is also designed to be FAIR, that is, it is findable, accessible, interoperable, and reusable. The latter two criteria are fulfilled by their creative commons licenses but also by their connection to other databases such as the Reptile Database (especially for names) and Vertnet (for specimen information, including localities). The Reptile Database also provides and/or directly links to the original description and other relevant literature so that access to the primary sources is provided.

### Morphological features

Each of our photos shows dozens or hundreds of individual scales and their features (size, shape, relationships, pattern, surface features etc), hence we cautiously estimate that we have documented at least a million, but probably millions of individual characters with our project. This database will become especially valuable when used in combination with other data sources such as the Reptile Database which now has descriptions of about 8000 species, mostly from the primary literature (Uetz et al. 2023). Most of these descriptions are of little use without images and many of the images in the primary literature are hard to find or behind paywalls (or there are no images, as in the majority of older descriptions). However, many historical descriptions have detailed descriptions of scalation features, and these will become available in conjunction with our image database. For instance, some lacertid genera have a single or paired postnasal and this character is easy to recognize, even for non-experts (Fig. 1). However, some characters are relative, such as having “enlarged masseteric scales”: without at least two photos to compare the sizes of scale in different specimens it is practically impossible to know what “enlarged” means, given that most lacertids have more or less enlarged masseteric scales (Fig. 1).

### Limitations

Obviously, this project is only a first step towards a complete database of reference images. Even if we had images of all 12,000 reptile species, that would still not reflect the diversity and variation within species, as geographic and individual variation needs to be represented by additional images. Similarly, even those specimens that are represented often lack critical meta-data, such as precise localities or biological data such as sex or age (the latter can be estimated from size information, but that is usually not included in collection data).

A final concern is the notorious problem of taxonomic uncertainty: during the process of taking pictures, we found several mis-identified specimens which may have been simply mis-identified but it is increasingly common that species change names because of splitting. That is, if a species gets split into multiple other species, a subset of specimens needs to be re-allocated to the newly split off species which rarely happens in collections. This problem can only be solved by taxonomists and collection managers who closely follow the literature (or online resources such as the Reptile Database) and relabel those specimens as needed. However, our online resource will also allow the community to re-examine specimens and thus correct such misidentifications.

While the reference image database could also incorporate images of live animals, these specimens are likely not available for follow-up studies.

### Linking data and databases

A critical component of this project was the linking of specimen data to both images but also collection databases such as VertNet. However, numerous other links are needed and possible, e.g., links to observation data (iNaturalist, GBIF), conservation data (IUCN), or DNA sequence databases which increasingly reference specimen information. DNA sequences will become increasingly important once phenotypes can be linked to genome data.

### Usage of reference images

The reference library has several possible uses. The first and most obvious is to use it in taxonomic studies for comparison and for the identification of species. This includes the use by amateurs, e.g., in iNaturalist where many photos do not show sufficient details for exact species identification. Second, reference images will allow researchers to investigate traits much more easily and systematically, especially when they are combined with ontologies of terms. For instance, lacertids such as the species shown in Figures 1 and 2 have essentially all the same types of scales. As long as a user knows which characters to look for, all images with that character can be easily found, simply by searching for all images that show, for instance, the dorsal side of a head (which has prefrontal, frontal, parietal, etc. scales). This could also facilitate the analysis of geographic variation. Third, we also envision possible uses for image analysis and automated analysis of the dataset. Reptiles are particularly suited for image analysis as the scales allow for easy demarcation and quantification, in contrast to the skin of amphibians or the fur of mammals which has much fewer landmarks. There are many more uses possible, and we invite the scientific community to explore such uses.

**Fig. 2.**
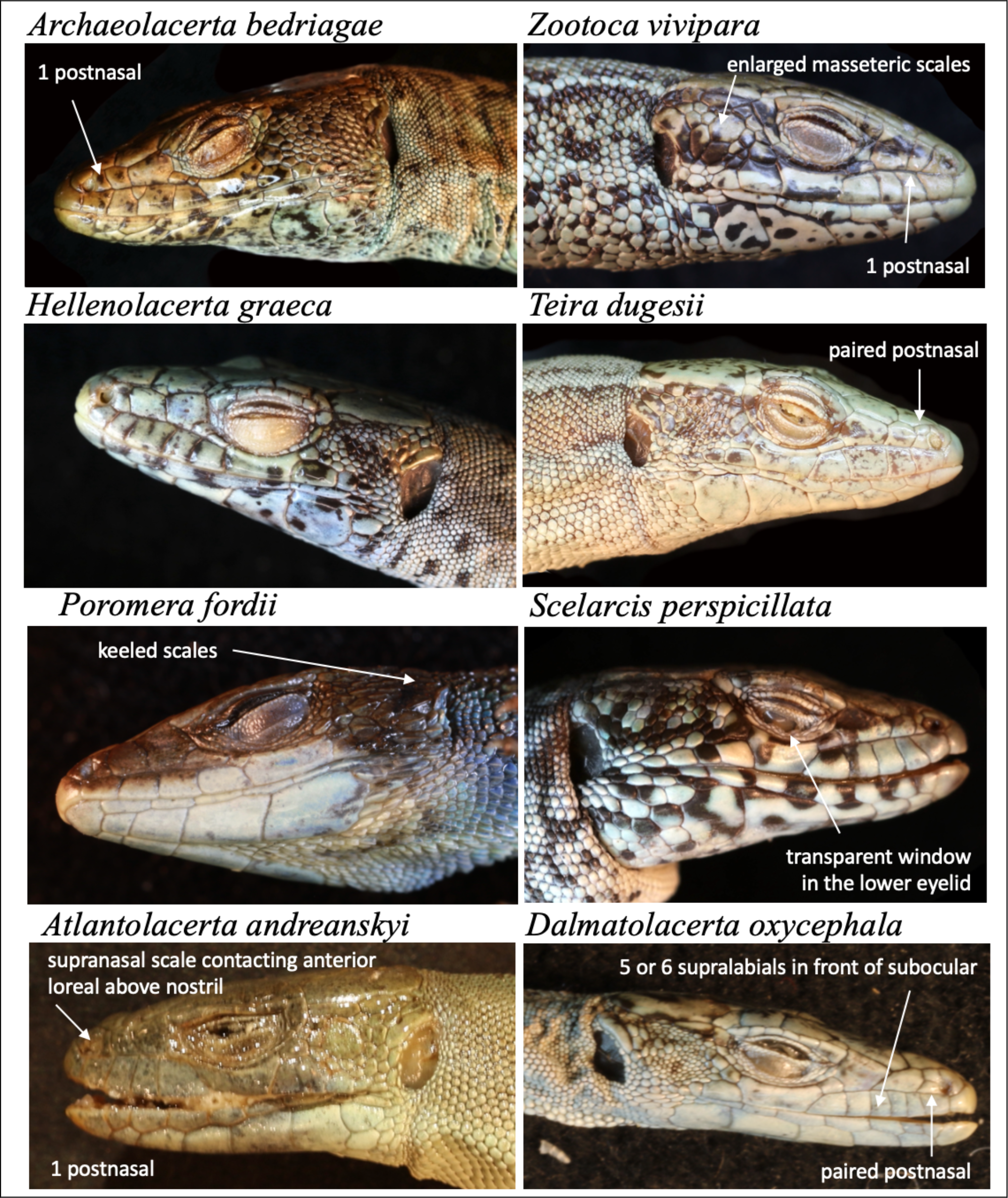
A comparison of 8 Lacertid genera using reference photos. Six species are in the subfamily Lacertinae, tribus Lacertini (*Archaeolacerta, Dalmatolacerta, Hellenolacerta, Scelarcis, Teira*, and *Zootoca*), but two are not in this tribe (*Atlantolacerta, Poromera*). Selected diagnostic traits are indicated.

### Future improvements and outlook

Most obviously, we need to expand the image collection to all reptile species, and eventually to all species in the animal and plant kingdoms. It remains to be seen to what extent this is possible, but with the inclusion of variation, the task as well as the possibilities are nearly endless. While it may be desirable to have 3D scans of all species, including photogrammetric images, the added benefit appears to be limited at this time, given the huge investment for 3D imaging, in addition to the storage and time requirements.

It will remain a challenge to maintain databases and the links to numerous other resources, but the increasing use of standardized identifiers (such as NCBI taxonIDs) should facilitate that in the future.

## Acknowledgments

We thank the collection managers and curators who provided access to the specimens in their care, namely Dane Trembath (AMS), Stephanie Tessier (CMNAR), Andreas Schmitz (MHNG), Nicolas Vidal (MNHN), Bryan Stuart (NCSM), Eduard Stöckli (NMBA), Jane Melville (NMV), Silke Schweiger and Georg Gassner (NMW), Andrew Amey (QM), Garin Cael (RMCA), Ralph Foster (SAMA), Gunther Köhler (SMF), Alexander Kupfer (SMNS), Shai Meiri (TAU), Coleman Sheehy (UF), Paul Doughty (WAM), Flecks Morris (ZFMK), Frank Tillack (ZMB), Chan Kin Onn (ZRC), and Michael Franzen (ZSM). Kenzley Adolphe and Tanya Berardini generously helped with this project’s implementation at Morphobank.

## Supplementary tables and data availability

**Supplementary Table S1**. List of all images, species, and specimens with their collection and catalog numbers, as well as the body parts shown in the image (Excel spreadsheet), available at http://dx.doi.org/10.6084/m9.figshare.25366564.

All images are available organized by collection at the following URLs (Figshare):

AMS http://dx.doi.org/10.6084/m9.figshare.25353253

CMNAR http://dx.doi.org/10.6084/m9.figshare.25353361

MHNG http://dx.doi.org/10.6084/m9.figshare.25353364

MNHN http://dx.doi.org/10.6084/m9.figshare.25353370

NCSM http://dx.doi.org/10.6084/m9.figshare.25354276

NMBA http://dx.doi.org/10.6084/m9.figshare.25355707

NMV http://dx.doi.org/10.6084/m9.figshare.25355722

NMW http://dx.doi.org/10.6084/m9.figshare.25355725

QM http://dx.doi.org/10.6084/m9.figshare.25355740

RMCA http://dx.doi.org/10.6084/m9.figshare.25355743

SAMA http://dx.doi.org/10.6084/m9.figshare.25355749

SMF http://dx.doi.org/10.6084/m9.figshare.25355752

SMNS http://dx.doi.org/10.6084/m9.figshare.25355755

TAU http://dx.doi.org/10.6084/m9.figshare.25355758

UF http://dx.doi.org/10.6084/m9.figshare.25355761

WAM http://dx.doi.org/10.6084/m9.figshare.25355767

ZFMK http://dx.doi.org/10.6084/m9.figshare.25355770

ZMB http://dx.doi.org/10.6084/m9.figshare.25355785

ZRC http://dx.doi.org/10.6084/m9.figshare.25355794

ZSM http://dx.doi.org/10.6084/m9.figshare.25355797

High resolution images are available from http://morphobank.org/permalink/?P5121.

